# Genome-wide association studies of polygenic risk score-derived phenotypes may lead to inflated false positive rates

**DOI:** 10.1101/2022.09.10.507410

**Authors:** Emil Uffelmann, Danielle Posthuma, Wouter J. Peyrot

**Affiliations:** Department of Complex Trait Genetics, Center for Neurogenomics and Cognitive Research, Amsterdam Neuroscience, Vrije Universiteit Amsterdam; Department of Child and Adolescent Psychiatry and Pediatric Psychology, Section Complex, Trait Genetics, Amsterdam Neuroscience, Vrije Universiteit Medical Center, Amsterdam University Medical Center, Amsterdam, The Netherlands; Department of Psychiatry, Amsterdam UMC, The Netherlands

## Abstract

In a recent study, a polygenic risk score (PRS) for Alzheimer’s disease was used to construct a new phenotype for a subsequent genome-wide association study (GWAS). Here we show that the applied method, in which the same genetic variants are used to construct the PRS-derived phenotype as well as to assess their effect in a GWAS of the same phenotype, leads to inflated false positive rates. We illustrate this bias by simulation. We first simulate an initial discovery cohort, and run a GWAS of a disorder like Alzheimer’s disease. We then simulate a target cohort, in which we construct a PRS based on the initial GWAS results. Following the published study, we select the bottom and top 5% of individuals in the PRS distribution and define them as controls and cases. Lastly, we run a GWAS on the new PRS-derived phenotype using all genetic variants. We show that at a significance threshold of 5 × 10^−8^, false positive rates are inflated up to 0.004 (an 80,000-fold increase compared to 5 × 10^−8^). We also show that such inflation can be prevented by excluding all variants that were used to construct the PRS (as well as all variants in linkage disequilibrium), when a GWAS on a PRS-derived phenotype is conducted.

## Main

Gouveia and colleagues (2022)^1^ conducted a genome-wide association study (GWAS) of a polygenic risk score (PRS)-derived phenotype (N = 37,784), in which they identified 246 independent loci and 473 lead SNPs. This is an enormous increase compared to the most recent and largest GWAS of AD^2^ (N = 1,126,563), which identified 38 loci. Here we show that the applied approach by Gouveia and colleagues may lead to an inflated false positive rate.

In this approach, beta-estimates from a recent GWAS of Alzheimer’s disease (AD)^3^ were used to construct PRSs in the European UK Biobank^4^ sample, using pruning and thresholding^5^ with a p-value threshold of 5%. Next, a new case-control phenotype was constructed based on the bottom and top 5% of the PRS distribution, removing 90% of their initial sample. Lastly, a GWAS was conducted on this new PRS-derived phenotype. The authors reasoned that by enriching the sample for individuals with known AD-associated variants, you may also enrich for unknown AD-associated variants. Our major concern is that the applied approach used the same single-nucleotide polymorphisms (SNPs) to construct, as well as to predict the phenotype. In other words, the phenotype was partly regressed on itself, which can inflate test statistics.

We performed simulations roughly emulating the approach (see Methods). In short, we simulated individual phenotypes under a liability threshold model and genotypes that loosely reflect the genetic architecture of AD^2,3,6^ (excluding the APOE locus) including 170,000 independent SNPs of which 1200 were causal and 168,800 were non-causal (null-SNPs). We then simulated a discovery sample such that the PRS explains approximately 5% of the phenotypic variance on the liability scale (N = 366,771). We ran a GWAS of AD in this discovery sample and used the estimated betas to construct a PRS in a target sample (N = 300,000). We then selected individuals in the top and bottom 5% of the PRS distribution (N = 30,000) and ran a second GWAS on this new PRS-derived case-control phenotype. The target cohort overlapped to varying degrees with the discovery cohort (i.e. 0%, 50%, and 100%), noting the AD GWAS summary statistics used by Gouveia and colleagues (2022)^1^ also contained the UK Biobank.

Our results show highly inflated false positive rates in the GWAS of the PRS-derived phenotype (see Fig. 1 and Supplementary Table). Across all null-SNPs and when there is no overlap between discovery and target cohort, the false positive rate was 0.0024 (s.e.m. = 1 × 10^−5^), which constitutes a 48,000-fold increase compared to a well-controlled false positive rate of 5 × 10^−8^ (see Supplementary Fig. 1 for α = 0.05). Another metric to quantify bias that is independent of the chosen significance threshold is the variance of test statistics, which should be 1 for a test with a well-calibrated false positive rate. Similarly, we find the mean variance of test statistics for all null-SNPs to be inflated at 1.64 (s.e.m. = 6 × 10^−4^) when there is no overlap between the target and discovery cohort. This inflation is driven by null-SNPs that were used to construct the PRS-derived phenotype. The variance of test statistics of these null-SNPs was equal to 13.8 (s.e.m. = 0.014) and the false positive rate was 0.05 (s.e.m. = 0.0003, a 1 × 10^6^-fold increase) when there was no overlap, while null-SNPs which were not used to construct the PRS-derived phenotype did not show any inflation. We also looked at the number of false positive associations per study (i.e. false positive rate times the number of null-SNPs considered), which was 408 in total when there was no overlap and was fully driven by SNPs used to construct the PRS-derived phenotype.

**Figure 1.**
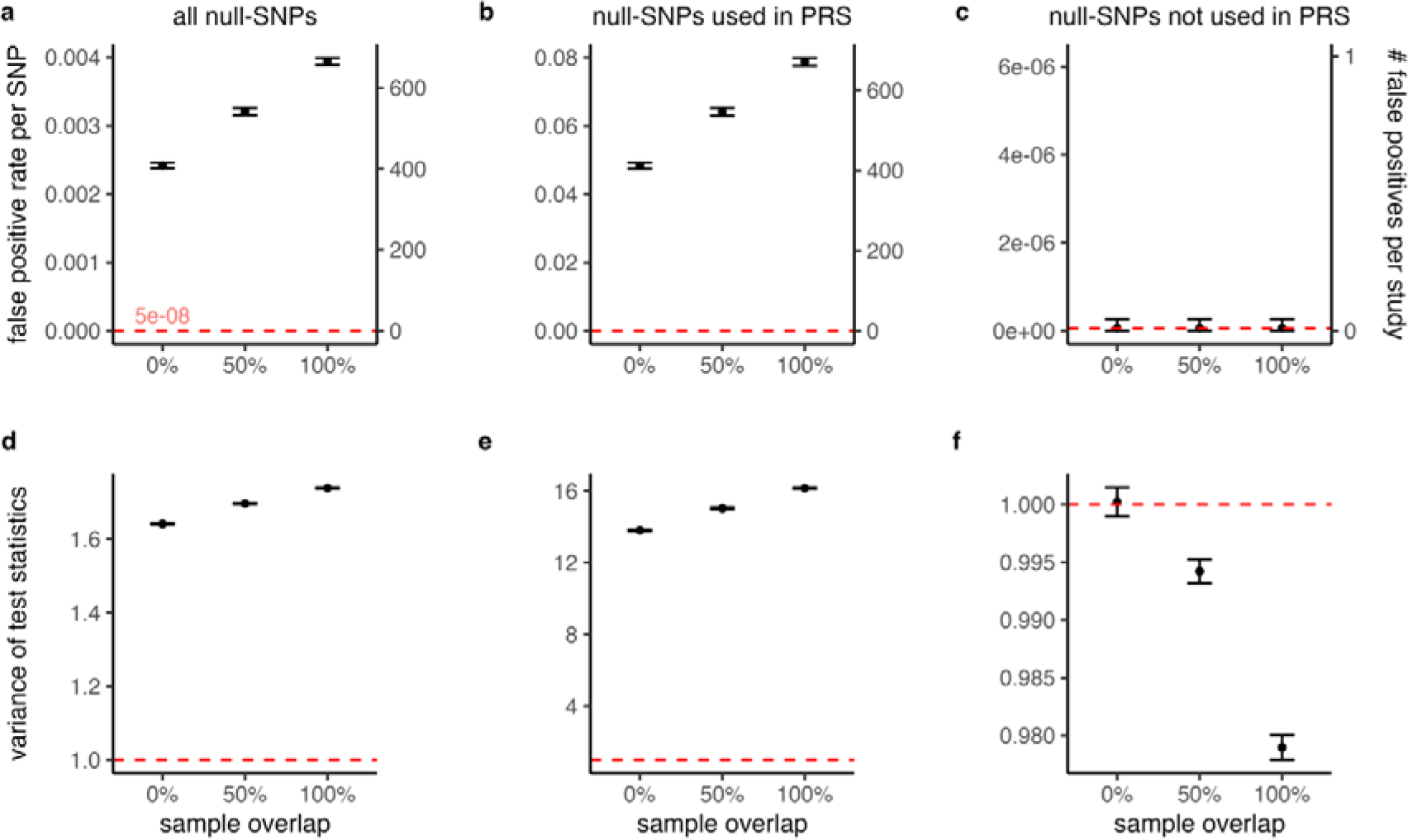
Inflated false positive rates and test statistics in GWAS of PRS-derived phenotype. The false positive rates **(a-c)** and variance of the test statistics **(d-f)** are displayed for varying degrees of sample overlap between discovery and target cohort in a GWAS of a PRS-derived phenotype. Across 100 simulation runs, we observe highly inflated false positive rates and test statistics. For all null-SNPs, the mean false positive rate ranges between 0.0024 (0% overlap) and 0.0039 (100% overlap) at a significance threshold of 5 × 10^−8^ **(a)**. Similarly, the mean variance of the test statistics ranges between 1.64 and 1.74, while it is expected to be 1 for a well-calibrated test (red line) **(d)**. Null-SNPs used to construct the PRS-derived phenotype show the highest inflation **(b, e)**, while all other null-SNPs do not show any inflation **(c, f)**. As such, the inflation in all null-SNPs is driven by SNPs used to construct the PRS-derived phenotype. Increasing overlap between the target and discovery cohort exacerbates the inflation. For null-SNPs not used to construct the PRS-derived phenotype, we observe a deflation of test statistics with increasing overlap **(f)**. We additionally plot the number of false positive associations per study (i.e. false positive rate per SNP times the number of SNPs considered). The mean number of false positives ranged between 408 and 665, and is driven by null-SNPs used to construct the PRS-derived phenotype. The error-bars show the 99.9%-confidence interval of the mean. See Supplementary Table for descriptive statistics.

Overlap between the discovery and target cohort exacerbated false positive rate inflation, increasing the false positive rate to 0.04 (s.e.m. = 1.6 × 10^−5^) across all null-SNPs when there was complete overlap. Similarly, the mean variance of test statistics increased to 1.74 (s.e.m. = 6 × 10^−4^) and the number of false positive associations increased to 665 (s.e.m. = 2.7). Interestingly, overlap between the discovery and target cohort inflated the test statistics for null-SNPs used to construct the PRS-derived phenotype but deflated it for all other null-SNPs. The reason for this is that p-values for null-SNPs will be correlated between the GWAS for AD and the GWAS for the PRS-derived phenotype when there is sample overlap (because AD and the PRS-derived phenotype are correlated and the same individuals are used). Selecting SNPs with p-values smaller than 0.05 for the PRS similarly selects SNPs not part of the PRS with p-values larger than 0.05. As a consequence, the GWAS of the PRS-derived phenotype will have deflated test statistics at null-SNPs not included in the PRS.

Next, we varied the p-value threshold for inclusion in the PRS (i.e. varying the threshold from 0.05 to 1 and 5 × 10^−8^, thus including either all SNPs or only genome-wide significant SNPs, respectively). We found that using all SNPs in constructing the PRS-derived phenotype reduced the inflation of false positive rates (as well as the variance of test statistics and number of false positives, see Supplementary Fig. 2). However, increasing the p-value threshold to 5 × 10^−8^ resulted in false positive rates and variances of test statistics that are not inflated. This is because almost no null-SNP had such a low p-value for AD, and thus almost no null-SNPs were used to construct the PRS-derived phenotype.

To summarize, Gouveia and colleagues (2022)^1^ used a new study design with the aim to improve the power for a GWAS of Alzheimer’s disease. Based on simulations, we showed that this approach may lead to inflated false positive rates of up to 80,000-fold increases. The reason for this is that the same SNPs used to construct the PRS-derived phenotype were subsequently tested for association with this newly constructed phenotype. We found the false positive rate inflation was more pronounced in the case of sample overlap between the discovery and target cohort. Our results show that false positive rates are not inflated when the GWAS of the PRS-derived phenotype is performed on SNPs that were not also used to construct the PRS. However, we note that when there is linkage disequilibrium between SNPs included in the PRS and null-SNPs not included in the PRS this could still result in an inflated false positive rate. An appealing approach may be to use a leave-one-chromosome-out approach, where the PRS is constructed using 21 chromosomes, and the GWAS of the PRS-derived phenotype only uses the 22nd left-out chromosome (repeated 22 times so that all chromosomes are left out once). Still, we note SNPs can also be correlated across chromosomes due to e.g. non-random mating^7^ which could in theory also lead to inflated false positive rates for this approach; however, we are not certain about the extent of this inflation which could well be negligible. See the Supplementary Note for a short discussion of some other approaches analyzing (partly) PRS-derived phenotypes^8,9^.

To conclude, phenotype definitions based on PRSs require careful consideration in subsequent GWAS. We recommend excluding any SNP (and those in linkage disequilibrium) from the GWAS that was used to construct the PRS-derived phenotype to prevent inflation of false positive rates.

## Methods

### Simulation

We simulated individual genotype and phenotype data based on the liability threshold model. Our chosen parameters were loosely based on Alzheimer’s disease^2,3,6^, with a population and sample prevalence of 5%, SNP-heritability (*h*^2^_SNP_) of 10% on the liability scale, and a PRS that explains 5% of the variance (*R*^2^) on the liability scale. We simulated a total of 170,000 SNPs in linkage equilibrium with a minimum minor allele frequency of 0.1%, as this was the number of pruned SNPs used by Gouveia et al (2022)^1^. Out of these, 1200 SNPs were causal, as previously estimated for Alzheimer’s disease^6^, and 168,800 were non-causal. We used the avengeme R package to calculate the number of individuals required for the discovery cohort to produce a PRS that explains the desired *R*^2^ value on the liability scale^10^. We simulated individuals and their liabilities, such that individuals with liabilities larger than the liability-threshold are designated cases, and otherwise controls. We repeatedly simulated individuals until we reached the desired number of individuals (N = 366,771 discovery, N = 300,000 target). We repeated the simulation for three target cohorts. That is, within the same simulation run, one target cohort was fully independent of the discovery cohort (0% sample overlap), in the other 50% (and 100%) of individuals were also present in the discovery cohort. Next, we ran a GWAS in the discovery cohort using plink version 1.9^11^. Using the estimated betas, we calculated PRS in the target cohorts to determine the top and bottom 5% of the PRS distribution to define the PRS extremes (i.e. the PRS-derived phenotype), and thus removed 90% of the sample. Lastly, we ran a second GWAS of the PRS-derived phenotype (N = 30,000) and recorded the false positive rate and the variance of test statistics. We repeated the simulation 100 times. We performed several model checks to ensure our simulations have the desired characteristics; specifically, we verified that the false positive rate and test statistics are not inflated for the primary GWAS of Alzheimer’s disease.

## Data and code availability

All code used for this manuscript is available at https://github.com/euffelmann/paper-ad_prs_extremes. Simulation results can be downloaded from https://doi.org/10.5281/zenodo.6818449.

## Acknowledgements

D.P. is supported by the Netherlands Organization for Scientific Research - Gravitation project ‘BRAINSCAPES: A Roadmap from Neurogenetics to Neurobiology’ (024.004.012) and the European Research Council advanced grant ‘From GWAS to Function’ (ERC-2018-ADG 834057). W.J.P is supported by a NWO Veni grant (91619152).

## Competing interests

The authors declare no competing interests.

## Author contributions

E.U. conceived the project, wrote the manuscript text and prepared the figures. W.J.P. and E.U. wrote the analysis code. W.J.P. and D.P. supervised the project. All authors discussed and commented on the manuscript.

## Notes

https://github.com/euffelmann/paper-ad_prs_extremes

### Competing Interest Statement

The authors have declared no competing interest.

