## Supplement for "Genome-wide association studies of polygenic risk score-derived phenotypes may lead to inflated false positive rates"

Affiliations:

### Supplementary Note

Polygenic risk scores (PRS) have been used in other ways to define new phenotypes or to improve power. For example, a recent study of resilience to schizophrenia^1^ also studied a PRS-derived phenotype but removed all schizophrenia risk SNPs, as well as SNPs in linkage disequilibrium with risk SNPs (*R*^2^ > 0.2), prior to running a GWAS. However, remaining linkage disequilibrium may still slightly inflate the false positive rate of the study. Other approaches utilize known SNP-associations to increase power by conditioning on PRS in combination with a leave-one-chromosome-out approach. The rationale behind this approach is that conditioning the phenotype on the PRS (e.g. based on chromosome 1-21) reduces the phenotype’s overall residual variance, thereby increasing the power for SNPs (on chromosome 22). While this approach should reliably improve power for continuous traits, such conditioning can reduce power for case-control traits in ascertained samples^2^, where independent predictors can become correlated due to collider bias. We note Zaitlen and colleagues (2012)^3^ addressed this issue and developed a method to improve power in GWAS by conditioning on known risk variants for case-control phenotypes based on the liability threshold model. In short, they transform a case-control phenotype to a quantitative liability phenotype, while taking the effect of known risk variants (or potentially other covariates such as a PRS) into account. They reported a well-controlled false positive rate, but observed that the power increase will be minimal for conditions that are not common (see Figure 2 in ref. ^3^), such as AD.

### Supplementary Table

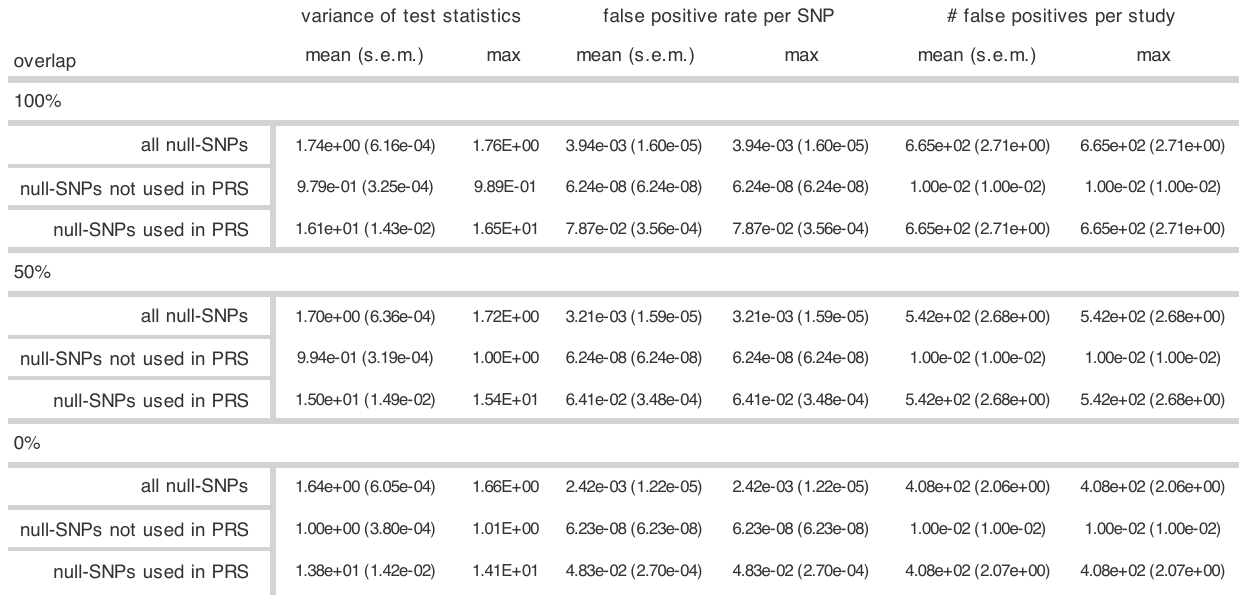

**Supplementary Table Descriptive statistics for GWAS of PRS-derived phenotype.** The values in this table are plotted in main Fig 1. The false positive rates, variances of the test statistics, and number of false positive associations are displayed for varying degrees of sample overlap between discovery and target cohort in a GWAS of a PRS-derived phenotype. Across 100 simulation runs, we observe highly inflated false positive rates and test statistics. For all null-SNPs, the mean false positive rate ranges between 0.0024 (0% overlap) and 0.0039 (100% overlap) at a significance threshold of 5 x 10^-8^. Similarly, the mean variance of the test statistics ranges between 1.64 and 1.74, while it is expected to be 1 for a well-calibrated test. Null-SNPs used to construct the PRS-derived phenotype show the highest inflation, while all other null-SNPs do not show any inflation. The mean number of false positives ranged between 408 and 665, and is driven by null-SNPs used to construct the PRS-derived phenotype. s.e.m. = standard error of the mean.

### Supplementary Figures

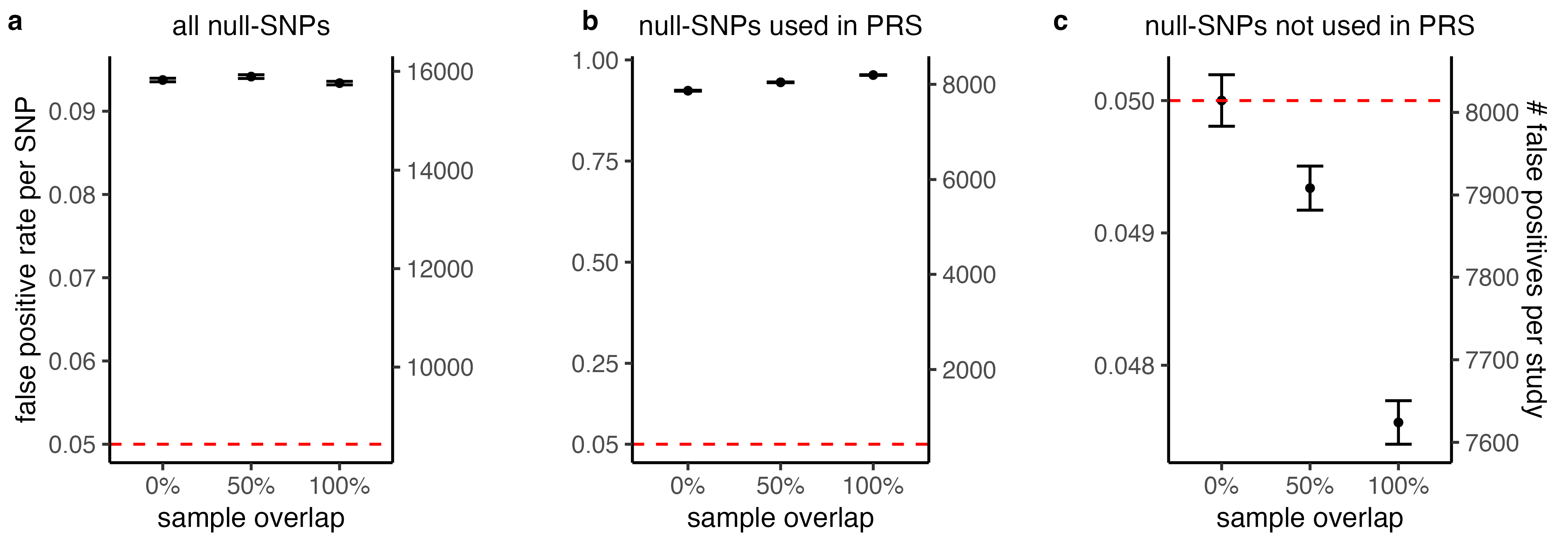

**Supplementary** **Figure 1 False positive rate (at α = 0.05) of GWAS of PRS-derived phenotype.** This figure is equivalent to main Fig. 1a-c, except that the significance threshold was set to 0.05 instead of 5e-8. The false positive rates are displayed for varying degrees of sample overlap between discovery and target cohort in a GWAS of a PRS-derived phenotype. Across 100 simulation runs, we observe highly inflated false positive rates. For all null-SNPs, the mean false positive rate ranges between 0.0934 and 0.0942, while it is expected to be 0.05 (red line) **(a)**. Null-SNPs used to construct the PRS-derived phenotype show the highest inflation **(b)**, while all other null-SNPs do not show any inflation **(c)**. As such, the inflation in all null-SNPs is driven by SNPs used to construct the PRS-derived phenotype. Increasing overlap between the target and discovery cohort exacerbates the inflation for null-SNPs used to construct the PRS-derived phenotype. For null-SNPs not used to construct the PRS-derived phenotype, we observe a deflation of false positive rate with increasing overlap **(c)**. We additionally plot the number of false positive associations per study (i.e. false positive rate per SNP times the number of SNPs considered). The error-bars show the 99.9%-confidence interval of the mean.

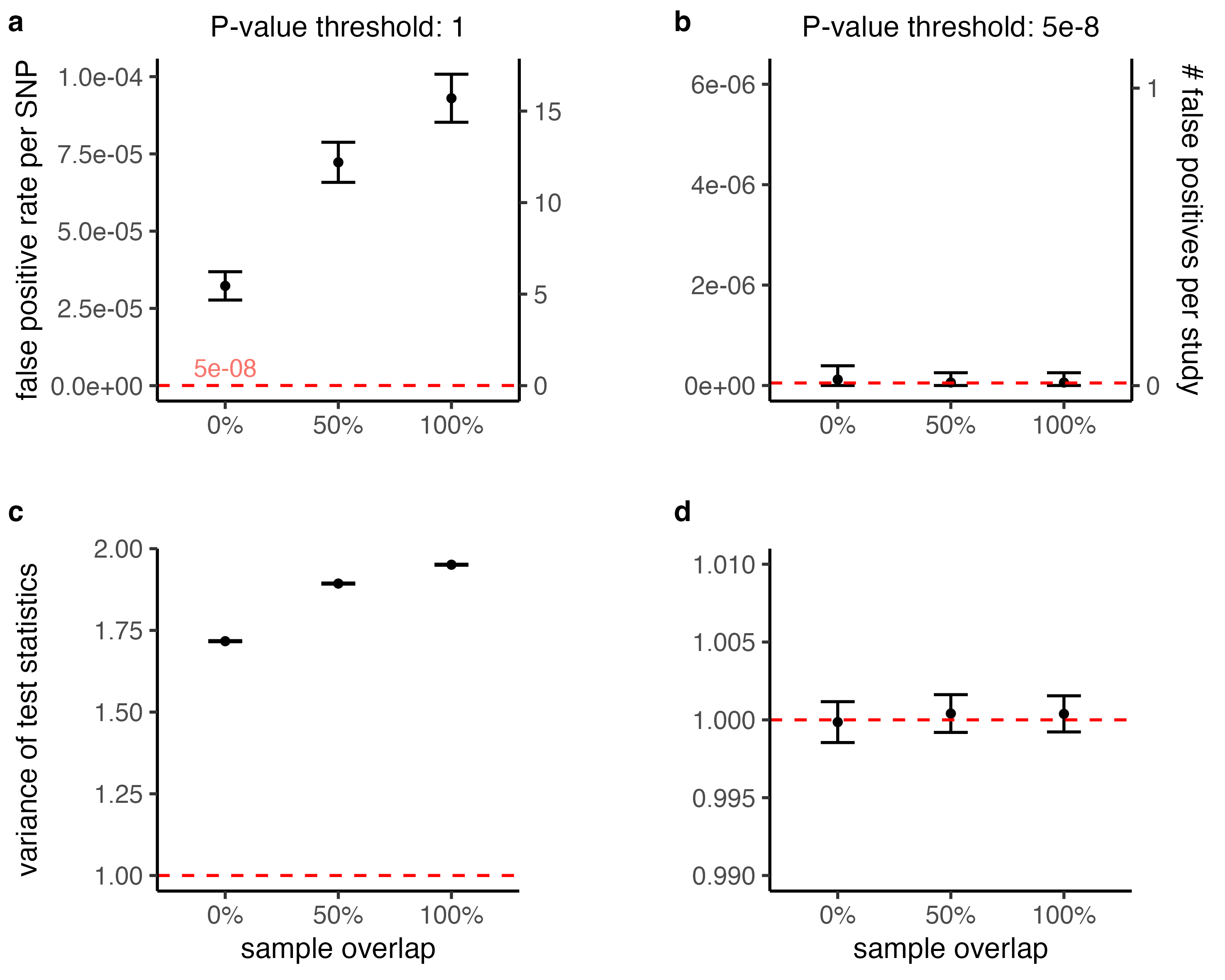

**Supplementary Figure 2 False positive rate at varying p-value thresholds for inclusion in the PRS.** This figure is equivalent to main Fig. 1a and d, except that all SNPs were used to construct the PRS-derived phenotype in **(a, c)** and only SNPs that are genome-wide significant were used to construct it in **(b, d)**. The false positive rates **(a, b)** and variance of the test statistics **(c, d)** are displayed for varying degrees of sample overlap between discovery and target cohort in a GWAS of a PRS-derived phenotype. In all panels the category “All null-SNPs” is shown. Note that in panel **(a, c)**, the previous categories “All null-SNPs” and “All null-SNPs in PRS” are equivalent. When using a p-value threshold of 1, the bias is diluted across all null-SNPs **(a, c)** and so the mean false positive rate, as well as the mean variance of the test statistics decreases. When using a p-value threshold of 5 x 10^-8^, almost no null-SNP is used to construct the PRS-derived phenotype **(b, d)**. Therefore, we observe no inflation. The error-bars show the 99.9%-confidence interval of the mean.
